# A System to Explore the Adaptive Dynamics of Multicopy Plasmids: The Role of Copy Number and Mutation Rate in Evolutionary Outcomes

**DOI:** 10.1101/2024.12.13.628408

**Authors:** Alexander (Olek) Pisera, Chang C. Liu

**Affiliations:** Department of Biomedical Engineering, University of California; Irvine, CA, 92617, USA; Center for Synthetic Biology, University of California; Irvine, CA, 92617, USA; Department of Chemistry, University of California; Irvine, CA, 92617, USA; Department of Molecular Biology & Biochemistry, University of California; Irvine, CA, 92617, USA

## Abstract

Multicopy plasmids are widespread in nature and compose a common strategy for spreading beneficial traits across microbes. However, the role of plasmids in supporting the evolution of encoded genes remains underexplored due to challenges in experimentally manipulating key parameters such as plasmid copy number and mutation rate. In this work, we developed a strategy for controlling copy number in the plasmid-based continuous evolution system, OrthoRep, and used our resulting capabilities to investigate the evolution of a conditionally essential gene under varying CN and mutation rate conditions. Our results show that low CN facilitates the faster enrichment of beneficial alleles while high CN promotes robustness through the maintenance of allelic diversity. High CN also slows the removal of deleterious mutations and increased the fraction of non-functional alleles that could hitchhike during evolution. This study highlights the nuanced relationships between plasmid CN, mutation rate, and evolutionary outcomes, providing insights into the adaptive dynamics of genes encoded on multicopy plasmids and nominating OrthoRep as a versatile tool for studying plasmid evolution.

## Introduction

Multicopy plasmids with copy numbers (CNs) up to hundreds per cell are widely distributed in nature. Such plasmids are found in bacteria^1^, yeast^2^, and mitochondria^2,3^, and confer traits to host cells that include antibiotic resistance^4^, anoxygenic photosynthesis^5^, synthesis of essential amino acids or vitamins^6^, and toxin production^7^. However, the widespread nature of multicopy plasmids presents a biological mystery^1^, because any given trait encoded on a multicopy plasmid can in principle be transferred to the single or low copy host genome where trait maintenance costs are minimized^8,9^. Furthermore, plasmid-encoded traits are subject to the cost of segregational drift where uneven plasmid segregation can lead to trait loss^10^.

The ubiquity of multicopy plasmids in the face of these clear costs has motivated extensive theoretical and experimental work to identify their evolutionary benefits. While the dominant hypothesis is that plasmids enable facile horizontal gene transfer compared to genomes^1^, thus increasing the accessibility of beneficial traits such as antibiotic resistance for potential hosts, recent work has proposed the additional possibility that plasmids may confer greater evolvability to encoded traits^11^. For example, multicopy plasmids increase the chance of sampling beneficial mutations by providing multiple copies of the target gene for mutation^12^, which should increase the rate of adaptation when mutation is limiting. It has also been suggested that multicopy systems enable “peak shifts” in the navigation of fitness landscapes wherein fit alleles in the same cell can conceal ones with deleterious mutations, allowing the latter to persist long enough to cross fitness valleys on the way to new fitness peaks^13^. In addition, multicopy plasmids may allow a cell to hedge against different environments by maintaining a diversity of alleles across different plasmid copies, thus preventing cells from over-specializing^11,14^ and securing high fitness in time-varying environments. However, these evolvability arguments for plasmids are tempered by the fact that in multicopy plasmid contexts, the contribution of beneficial alleles can be masked or diluted by other copies encoding other alleles. Therefore, deleterious mutations can persist, and beneficial alleles may be slow to enrich^14^ and quick to lose once found^10^.

While such complex cost-benefit tradeoffs underlying the evolvability of multicopy plasmid-encoded traits have been extensively modeled^14–16^, experimental efforts have proven challenging, in large part because it is difficult to control key variables such as CN and mutation rate without also changing other parameters such as plasmid type and stability. Additionally, experimental plasmid evolution studies typically rely on the low per base mutation rates of cellular DNA replication^10–12,17^, which can be too low to reliably sample beneficial mutations in plasmid-encoded genes even at high CN. Furthermore, such studies are often confounded by genomic adaptations^10,17^, since the genome shares the same per base mutation rate as the plasmid. This confound makes it difficult to observe the isolated contribution of plasmid evolution.

To overcome these challenges, we offer OrthoRep as an experimental multicopy plasmid evolution system wherein CN can be tuned, and mutation rate can be elevated for only the plasmid. OrthoRep was developed as a continuous directed evolution platform wherein a linear plasmid (p1) is replicated exclusively by a dedicated error-prone DNA polymerase (DNAP) that does not replicate genomic DNA^18^. Previous efforts on OrthoRep have yielded a collection of DNAPs spanning a range of error rates^19,20^ that afford the ability to control the frequency with which beneficial (and deleterious) mutations are sampled independent of the genome. In the current work, we add the ability to control p1’s CN independent of its per base mutation rate by tuning the expression level of the OrthoRep DNAPs that replicate p1. The result is a platform well-suited for investigating the effects of plasmid CN and mutation rate on evolutionary dynamics and outcomes.

We used our platform to study aspects of plasmid evolution by encoding the *Aspergillus nidulans amdS* amidase (amdSYM) of the formidase/acetamidase class^21^ on p1 under different CNs and mutation rates in the yeast, *Saccharomyces cerevisiae*. AmdSYM cleaves acetamide and related amides to release ammonia and has been previously used as a selectable marker that enables *S. cerevisiae* to metabolize acetamide as a sole nitrogen source^22^. In comparative evolution experiments, we evolved p1-encoded amdSYM to improve its ability to use butyramide instead of acetamide as the sole nitrogen source. The results of these experiments provided evidence for several predictions on how multicopy plasmids affect evolvability^10,11,23^. A key finding of our study is that evolution in low CN settings resulted in a higher fraction of improved alleles than did evolution in high CN settings. A related finding is that this higher fraction of improved alleles also had higher average fitness. This result can be explained by the fact that low CN means fewer copies are available to contribute to fitness and thus, there is more pressure on individual copies to perform well in contrast to the high CN case. Another key finding was that high CN supported the evolution of robustness to multiple environments by maintaining different alleles through “heteroplasmy”^11,14^. This was evidenced by our demonstration that compared to the low CN conditions, sets of evolved alleles from high CN experiments had greater ability to metabolize amdSYM’s original acetamide substrate when normalized to their ability to metabolize butyramide. We also found that when high mutation rates were used, especially in combination with high CN, more non-functional alleles were generated and maintained through masking by functional copies in the same cell. This was unsurprising since random mutations are generally deleterious^24^ so higher mutation rate should generate hypomorphic or non-functional alleles more rapidly – especially because in this experiment, wild-type (WT) amdSYM activity for butyramide, while suboptimal, was already high, leaving modest room for improvement. What was surprising, however, was the fact that we observed a bifurcation in allele fitnesses where at the end of evolution experiments, there was always a low mode of non-functional alleles and a high mode of functional sequences rather than a single mode somewhere in between. This finding adds a nuance to the evolutionary dynamics of multicopy plasmid evolution: even if CN stays constant across evolution, the effective CN on which selection is acting (the high mode of functional sequences) is dynamic. This offers a mechanism by which evolution can modulate the CN on which it is acting without changing the mechanisms of CN control. We discuss the implications of these findings in understanding the evolutionary dynamics of multicopy plasmids as well as the practicalities of carrying out directed evolution experiments using plasmid-based continuous evolution systems^25–27^, such as OrthoRep. Finally, we discuss the limitations of our studies to guide interpretation of our results and to motivate future work. Overall, we suggest that the level of CN and mutation rate control possible with OrthoRep makes it a versatile experimental platform to probe the evolutionary dynamics of plasmid-encoded traits.

## Results

### p1 copy number control

We hypothesized that the amount of DNAP available to replicate p1 would affect its CN, providing a straightforward method for CN control. To test this hypothesis, we modulated the expression level of OrthoRep’s error-prone DNAPs, synthetically encoded in the host yeast genome, using a set of characterized promoters spanning a wide range of strengths^28^. As media conditions are known to change the expression strength of promoters across the genome, we prioritized a set of promoters previously reported to have similar strengths in media with and without supplemental amino acids^28^. In addition, the *SAC6* promoter (pSAC6) was also included since we have used it previously for expressing OrthoRep error-prone DNAPs.

Thirty strains representing ten different promoters driving three recently published error-prone DNAPs – SgtKis, Trixy, and BadBoy3^20^ – were constructed and p1 plasmids encoding amdSYM and the LEU2 selection marker were transferred into these strains. Promoters reported to be weaker were generally associated with slower colony formation in media lacking leucine (Fig. S1), which provided a first indication that CN was being controlled by the availability of the DNAP under the reasoning that lower CN would result in less Leu2 protein expression. Based on this, one shared promoter set of three strains exhibiting fast growth was eliminated, as there was another set of equally fast-growing strains. An additional shared promoter set of three strains was also eliminated for practicality due to extremely slow growth.

For the remaining 24 strains representing 8 promoters, each driving the expression of three error-prone DNAPs at different strengths, we picked two colonies (biological replicates) into synthetic complete (SC) media lacking appropriate amino acids for selection and cultured for two days. We then split the cultures into two conditions, one with media approximating the eventual media for evolution experiments (*i.e*., synthetic complete (SC) with no supplemental amino acids, as nitrogen source selection would later be used for amdSYM activity) and another with SC lacking only the appropriate amino acids for strain and plasmid selection (*i.e*., SC without histidine, leucine, uracil, tryptophan, methionine, and cysteine (SC-HLUWMC)) to compare CN in both medias.

The CN of p1 was measured with qPCR for all strains, replicates, and conditions. Satisfyingly, the rank order of the CNs largely tracked the strength of the promoters driving the DNAP (Fig. 1). Per given polymerase, the CNs spanned an approximately 10-fold range. Per given promoter, the CN was generally inversely related with the error rate of the three DNAPs, SgtKis (2.4E-5), Trixy (4E-5), and BadBoy3 (1E-4) (Table S1). This inverse relationship was previously observed for OrthoRep DNAPs and likely reflects the way the DNAPs were engineered^19^ and/or fundamental fidelity/processivity relationships^29^. Interestingly, the CN of p1s replicated by the low error rate WT OrthoRep DNAP was not modulated by the range of promoter strengths tested, for reasons that we do not understand (Fig. S2A). For the media conditions lacking supplemental AA, the p1 CN was lower for most strains. Most dramatically, the p1 CN in strains where the DNAP was expressed from the legacy promoter, pSAC6, were reduced 2.5-fold in a single passage by changing to the media condition lacking supplemental amino acids (Fig. S2B). Taken together, these data validate our simple strategy for controlling the CN of p1 in OrthoRep.

**Figure 1.**
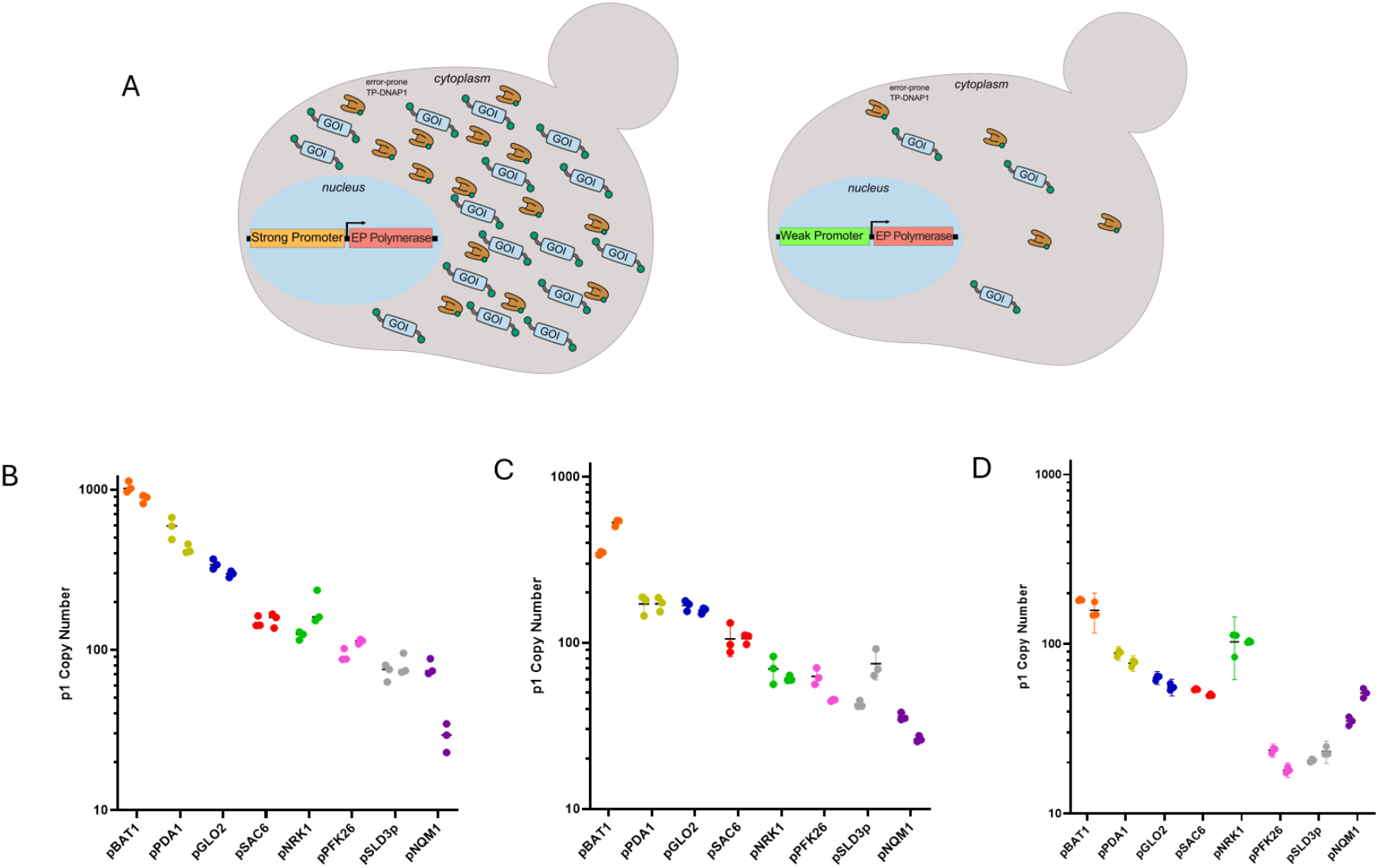
p1 CN achieved by 3 OrthoRep DNAPs expressed from 8 promoters. A) p1 CN is directly controllable by modulating the expression level of the error-prone OrthoRep DNAP. B) CNs achieved by SgtKis and different promoters. C) CNs achieved by Trixy and different promoters. D) CNs achieved by BadBoy3 and different promoters. GOI = gene of interest.

### CN is positively correlated with both expression strength and initial growth rates

Changes in CN results in changes in gene dosage, which should change the level of gene expression. However, the relationship between gene dosage and expression is not necessarily linear. To characterize this relationship for OrthoRep, we encoded mKate on p1 and measured fluorescence for strains maintaining p1 at various CNs using all three error-prone DNAPs. As shown in Fig. 2A, expression of mKate was positively and linearly correlated with CN across the range measured, suggesting that gene dosage through p1 CN acts as a simple multiplier of expression.

**Figure 2.**
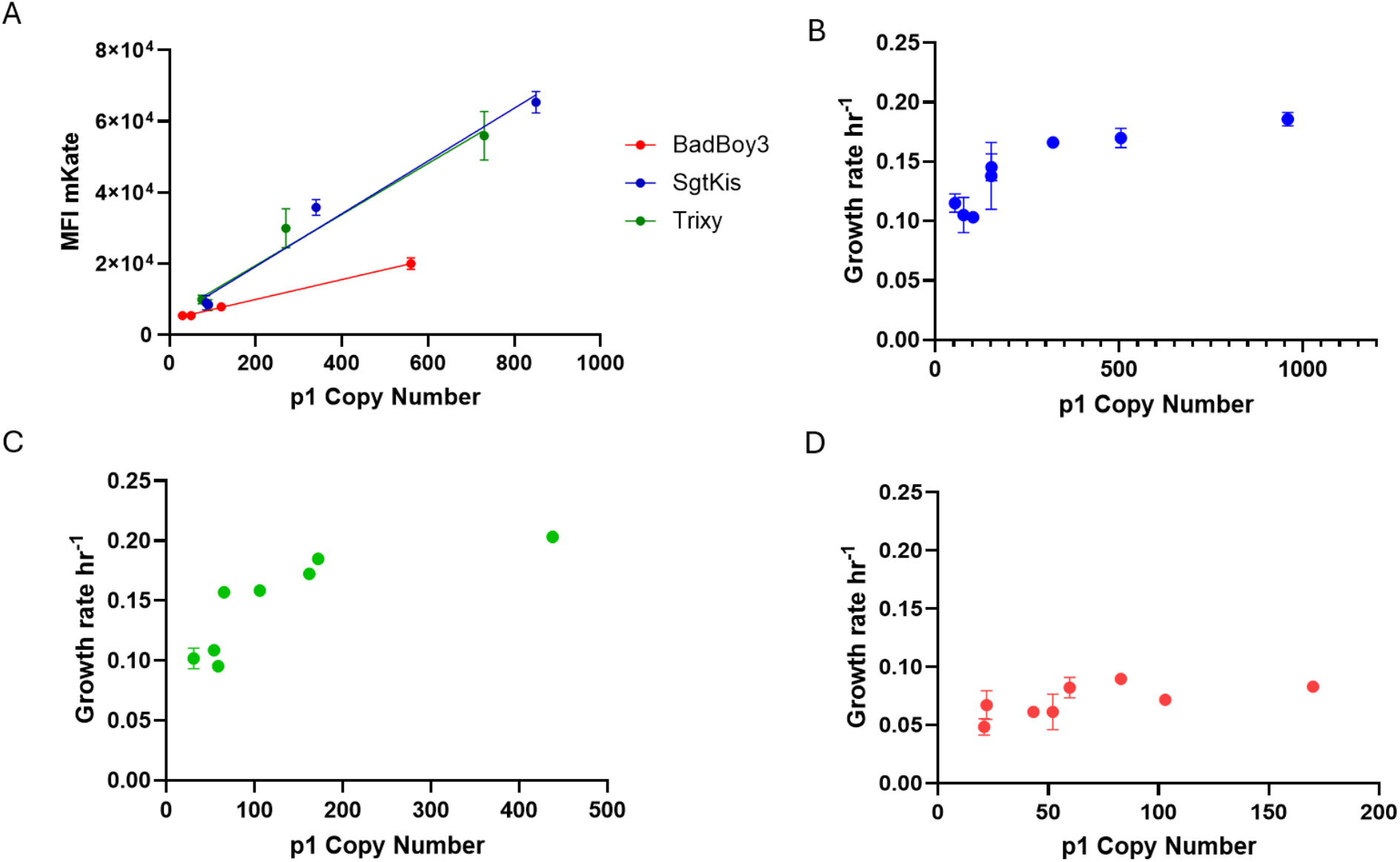
Influence of CN on expression and growth rate. A) Expression of a p1-encoded gene (mKate) is linearly correlated with CN, supporting a simple gene dosage model for total expression per cell. B-D) Growth rate is initially correlated with p1’s CN under selection for a p1-encoded gene but exhibits a sublinear relationship at high CN. All samples for SgtKis (B), Trixy (C), and BadBoy3 (D) were run in duplicate and the mean is plotted. Growth rate is reported as ln(2)/doubling time for the exponential growth rate constant. Error bars indicate range (the error bars are not visible on many samples due to close agreement between some duplicates). MFI = Mean Fluorescence Intensity

We also reasoned that by changing the expression level of the selection marker (*e.g*., *LEU2*) used to maintain p1, p1’s CN could influence the growth rate of cells. For the two error-prone DNAPs with lower error rates, SgtKis and Trixy, growth rate increased with increasing CN but began to plateau at a CN of 200, above which further CN increases hardly increased growth rate (Fig. 2B,C). The existence of a growth rate plateau is notable in its illustration that the relationship between an essential gene’s expression level and growth rate is non-linear: there is a point above which additional expression does not increase fitness. This will be important in explaining the dynamics of our evolution experiments with high CN systems (see Discussion). For BadBoy3, the growth rate was lower across all CNs and the correlation between CN and growth rate was less clear (Fig. 2D).

### Evolution of a p1-encoded gene

In order to study evolutionary dynamics as a function of CN and mutation rate, we encoded amdSYM on p1 in 8 strains per error-prone DNAP. Twenty-four total strains were evolved in duplicate whereby cells were passaged in minimal media containing a limiting concentration of butyramide (4 mM in the first 50 generations and then 2 mM in the remaining 50 generations) as the sole nitrogen source (Fig. 3A). For each passage, the optical density at 600nm light (OD_600_) was measured and normalized such that for each passage, the population sizes were similar when seeding the next passage. This also allowed for monitoring of the growth rate throughout the campaign.

**Figure 3.**
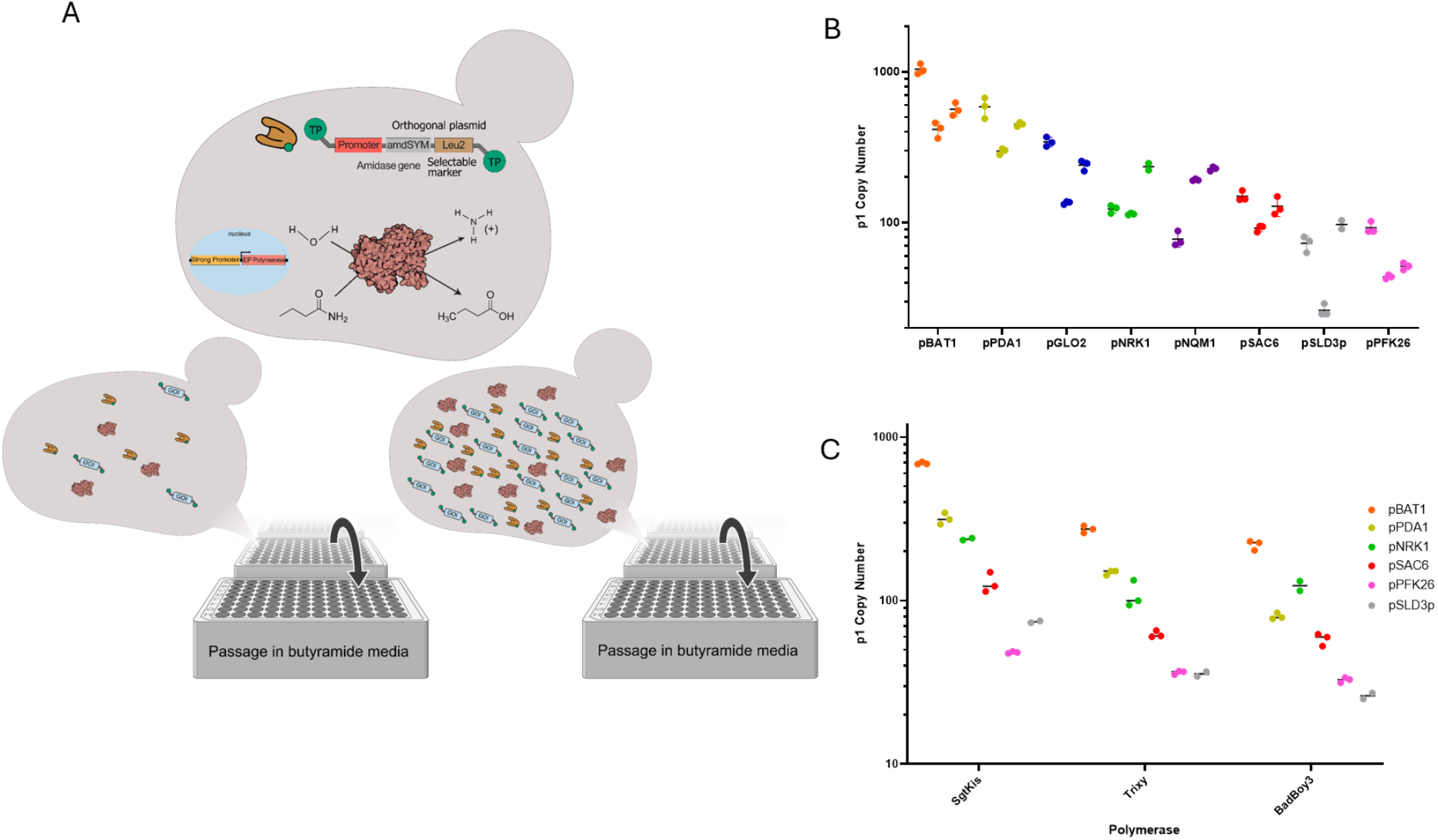
AmdSYM evolution experiment and CN stability. A) Evolution campaigns were carried out by passaging yeast strains with p1-encoded amdSYM in media with butyramide as the sole nitrogen source where butyramide was added at limiting concentrations (4 mM for the first 50 generations and 2 mM for the second 50 generations) to promote amdSYM adaptation. B) The CN of p1 during the evolution experiment remained mostly stable over ∼100 generations of selection for butyramide usage. Shown is the CN of one biological replicate of evolution with 8 promoters driving SgtKis measured at 3 timepoints, corresponding to passage 1, passage 7, and passage 13. C) For all 3 error-prone OrthoRep DNAPs, a wide range of CNs was maintained to the endpoint of the evolution campaign, representing a ∼10-fold CN range. These endpoints of evolution were used for further characterization.

During the evolution experiment, growth rate either quickly reached a plateau or started at the plateau such that low CN Trixy and SgtKis strains evolved for nearly an identical overall number of generations as the high CN populations, reflecting the same number of passages (Fig. S3A). For BadBoy3, the growth rate impact of reduced CN persisted throughout the evolution campaign (Fig. S3A). Notably, the growth rates for high CN strains were similar at the beginning and end of the evolution campaign, suggesting that amdSYM at high gene dosage was sufficient to support propagation at 4 mM butyramide conditions from the outset. The evolution campaigns were then carried out for approximately 100 generations for all populations. As a point of reference, at the error rates of the SgtKis (2.4E-5), Trixy (4E-5), and BadBoy3 (1E-4) DNAPs, this translates to the accumulation of roughly 3.8, 6, and 13 mutations per amdSYM sequence, respectively, under the null model of evolution.

We measured CN at the beginning, middle, and end of the evolution experiment. Even though pressure to increase amdSYM activity could have led to increases in CN especially for the low CN strains, CN stayed stable throughout the experiment (Fig. 3B). This indicates that CN control is forceful and OrthoRep is a suitable model system for isolating the effects of CN on evolution.

Our next task was to study the evolved alleles produced from the evolution experiments to understand the effect of CN and mutation rate on evolutionary outcomes and dynamics. For practical considerations that make it difficult to characterize evolved alleles from all 48 evolution experiments (24 evolution campaigns carried out in duplicate), we chose to focus on 18, representing a common set of 6 promoters for each error-prone DNAP, spanning a wide CN range (Fig. 3C).

### Outcomes of evolution

To characterize the results of the evolution campaigns, we subcloned the evolved alleles from p1 into a genomic integration cassette and included unique molecular identifiers (UMIs). This was achieved by amplifying the amdSYM coding region from p1 using PCR primers that added a promoter to drive genomic expression along with UMIs and flanks for efficient integration into the genome of yeast (Fig. 4A). The purpose of this subcloning was threefold. First, it would prevent further mutation of alleles during characterization. Second, it would downsample the number of mutant sequences into a practical range (∼10,000 sequences per evolution) for Nanopore sequencing to accurately match UMI to amdSYM mutant identity. Third, it would allow for high-throughput competition against a WT version of amdSYM (integrated in the same format with a known UMI) to measure the relative fitnesses of thousands of variants in parallel. Since each cell would have precisely one amdSYM variant encoded on its genome, we could compare the performance of outcomes, evolved in a multicopy context, at single allele resolution. Together, this assay allowed us to measure what fraction of sequences were functional (Fig. 4B), and gain insight into activity levels and their relationship to the enriched sequences (Fig. 4C-J).

**Figure 4.**
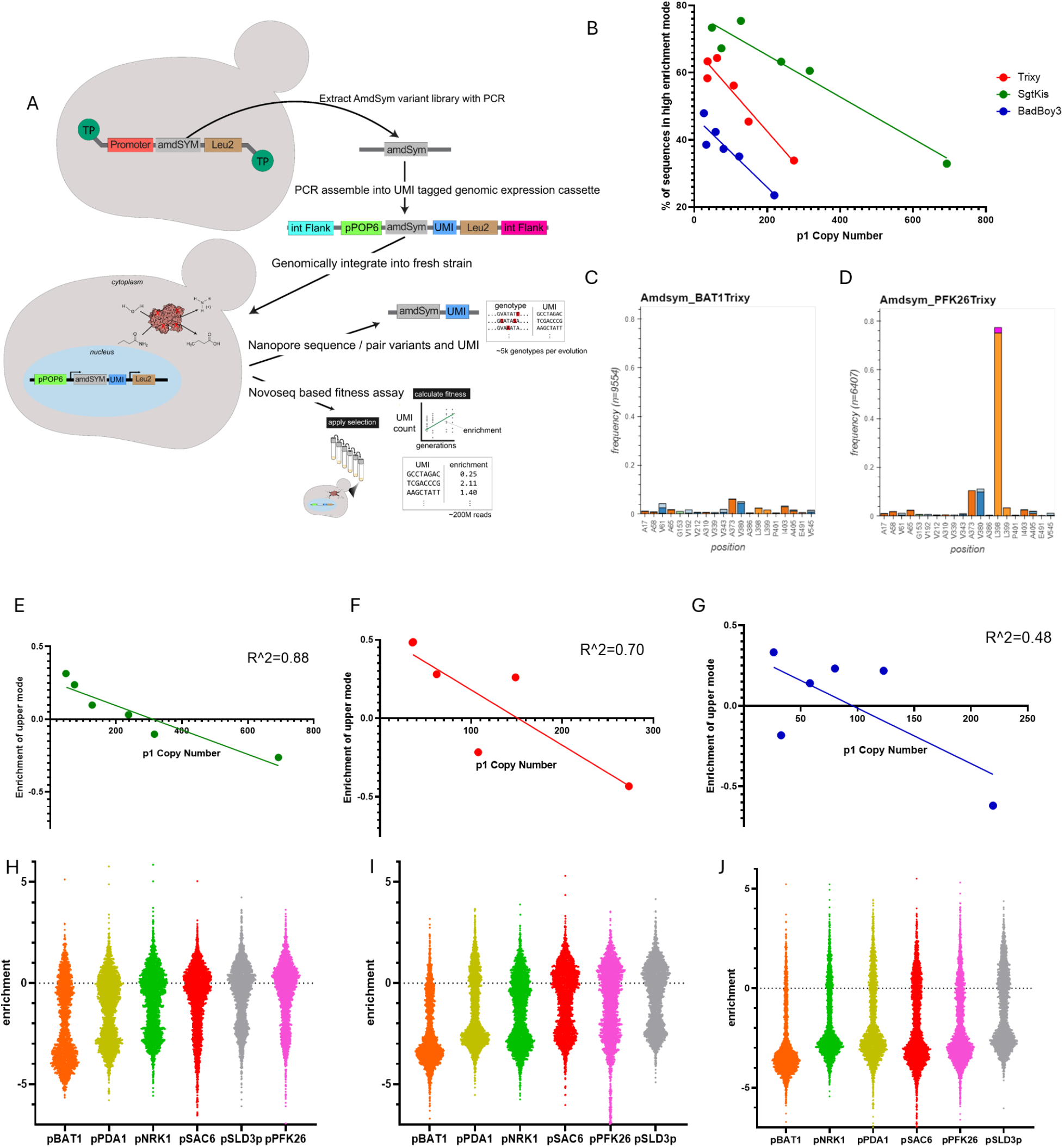
Assessment of AmdSYM variants produced by evolution under varying CN and error rate. A) Schematic describing the use of our in-house nanopore sequencing pipeline and Illumina enrichment assay. Briefly, variant libraries were assembled into a genomic integration cassette and transformed into competent yeast. The resulting variant libraries were sequenced and analyzed with nanopore to pair UMI with variant, and the same libraries were taken into a fitness assay to score each variant’s activity. B) The fraction of variants that were functional from each evolution condition plotted by CN at the final timepoint of evolution. C-D) Frequencies of sequences that contained each indicated mutation for representative outputs from a high CN evolution condition (C) and a low CN evolution condition (D). E-G) Analysis of the upper mode enrichment scores as a function of CN during evolution for SgtKis (E), Trixy (F), and BadBoy3 (G). The histograms used to generate this can be found in Fig. S5. The upper mode enrichment scores are negatively correlated with CN. H-J) The enrichment scores of 3000 random individual alleles evolved at different CNs (high to low from pBAT1 to pPFK26) for SgtKis (H), Trixy (I), and BadBoy3 (J). Enrichment is defined as log_2_(Ratiof_inal_/Ratio_start_) where Ratio is frequency relative to WT frequency. The WT only control, measured separately, grew 7.9 generations in this assay.

From the sequences of the alleles alone, a few key observations were made. First, the average number of nucleotide mutations per sequence did not vary significantly across CNs for any given DNAP (Fig. S3B), consistent with the expectation that our strategy of CN control did not impact the error rate, as it relies on changing DNAP expression levels but not their identity. Second, the fraction of sequences containing stop codons increased linearly with the CN, providing evidence of masking by high CNs where higher CN allowed more non-functional amdSYM mutants to hitchhike in the same cell as adapted functional amdSYMs (Fig. S3C). Third, different fractions of sequences containing a mutation known to increase amdSYM’s activity for butyramide (L398S) was also observed (Fig. S4A-E), which could be used as an indication of different degrees of amdSYM adaptation from different conditions. For example, the L398S mutation was present in 0.7% and 2% of sequences in the highest CN SgtKis and Trixy outcome alleles, respectively, whereas it reached 54% and 75%, respectively, in the lowest CN outcome alleles for the same DNAPs (Fig. 4C,D and Fig. S4A,B). This suggests that low CN can more quickly enrich a key beneficial mutation. The bulk growth of the pooled libraries in the competition assay reflected the inferred average fitnesses of all sequences, as the libraries produced by lower CN evolutions grew faster (Fig. S4F).

### Distribution of evolved allele fitnesses

To gain greater insights on evolutionary outcomes, we aimed to measure their distribution of fitnesses. We carried out a pooled competition experiment where the relative growth rates of clones expressing different amdSYM alleles could be determined in high throughput. Populations containing genomically-integrated amdSYMs were mixed at a known ratio with a reference strain encoding WT amdSYM. UMIs distinguished each variant from each other and the WT amdSYM. The populations were then grown in duplicate in media with butyramide (2 mM) as the sole nitrogen source. Genomic DNA from the initial population and the population after one passage, representing ∼7-8 generations of the WT amdSYM clone’s growth, was extracted and the UMIs were sequenced using the NovoSeq X sequencing platform. The enrichment of each UMI relative to WT over the total period of growth was determined for each variant, the log_2_ of which we defined as an enrichment score. Enrichment scores for the same variant between replicates were well correlated (R^2^ = 0.8) (Fig. S5A-C).

We noticed that for all conditions, the resulting distributions of enrichment scores were bimodal. We interpret the low mode to be composed of non-functional amdSYMs that were generated and maintained through masking by functional copies in the same cell during the evolution experiment. This is consistent with the fact that the distribution of enrichment scores for sequences containing stop codons completely overlapped with this low mode for all samples (Fig. S6A,B). As expected, non-functional sequences (*i.e*., the low mode) occupied a larger fraction of the overall distribution in the high CN (*versus* low CN) conditions, which we attribute to the fact that high CN can mask a larger number of hitchhiking non-functional alleles per cell during evolution. Also as expected, non-functional alleles were responsible for a larger fraction of the total distribution for higher error rate DNAPs (*e.g*., BadBoy3) compared to lower error rate DNAPs (*e.g*., SgtKis and Trixy). This is because BadBoy3’s higher error rate and different mutational biases^20^ should mean that it generates nonfunctional alleles more quickly than SgtKis and Trixy, since random mutations are mostly deleterious and transversion mutations are more non-conservative. For BadBoy3, 47% of evolved sequences were in the high enrichment score mode for the lowest CN evolution experiment while this percentage was only 23% for the highest CN evolution experiment (Fig. 4B). For SgtKis and Trixy, the high enrichment score mode accounted for 75% and 65% of the distribution, respectively, for the lowest CN evolution experiment outcomes in contrast to 33% and 34%, respectively, for the highest CN evolution experiment outcomes (Fig. 4B). Overall, this confirms that high CN slows the removal of low fitness alleles by purifying selection^10,14^.

If high CN can hide the low fitness of nonfunctional alleles from purifying selection, does it also hide the high fitness of beneficial alleles from positive selection? The trends described above for the beneficial L398S single mutation, which is generally more frequent in low CN versus high CN conditions, suggests that the answer to this question is yes. However, L398S is only a single known beneficial mutation. There may be other beneficial mutations and mutation combinations beyond (and even excluding) L398S at play. We therefore analyzed the enrichment scores of the high mode (*i.e*., functional sequences) across CNs. Although the differences aren’t dramatic, this analysis showed that the average enrichment scores of the high modes were indeed higher for low CN versus high CN evolutionary outcomes (Fig. 4F-H, Fig. S5). Our model for this relationship is that high CN resulted in reduced pressure for each individual copy of the amdSYM gene to improve during evolution, because there were a greater number of other functional copies contributing to fitness.

### Robustness of evolutionary outcomes in the ancestral environment

Evolution of amdSYM for improved butyramide usage under lower CN enriched beneficial variants to a greater degree than evolution under high CN on average. Alleles evolved under low CN could therefore be more specialized to butyramide and also compose a population with lower overall functional sequence diversity. This could lead to less amdSYM activity for amide nitrogen sources besides butyramide, including the ancestral acetamide substrate. In fact, we have measured such specificity tradeoffs for amdSYM harboring only the L398S mutation (Fig. S7A), which is a mutation we commonly identified in our evolved amdSYMs. To test this tradeoff idea, the growth rates of all CN varied populations for all three DNAPs were characterized in butyramide versus acetamide. The populations used were the same populations from the pooled competition experiments above wherein amdSYM alleles that resulted from different evolution experiments were genomically integrated into strains. However, unlike the pooled competition experiments, we first depleted populations of nonfunctional variants prior to growth rate characterization by first growing cells for one passage in limiting ammonia, then one passage in luxury concentrations (60 mM) of butyramide. This concentration is high enough that it should de-enrich nonfunctional amdSYM variants but not significantly enrich functional variants. The resulting populations were then grown in limiting concentrations of butyramide or acetamide as sole nitrogen sources. While the CN varying evolution experiments also have varying levels of expression per cell due to different functional CNs (Fig.S7B), we aimed to measure the relative activity per sequence in this assay (Fig. S7C,D).

After 48 hours of growth, we observed a clear trend for all three DNAPs where the functional allele sets from the high CN evolution conditions exceeded those from the low CN evolution conditions in their preference for acetamide as the sole nitrogen source (Fig. 5). This is consistent with a tradeoff model for the type of evolutionary outcomes evolved under high versus low CN. On the one hand, the activity of individual alleles may on average be better adapted for the target evolutionary pressure (butyramide usage) when adaptation occurs under low CN. However, the fact that high CN effectively acts to reduce the selection pressure on individual copies during evolution – thus preventing over-specialization of individual alleles and/or allowing more diverse alleles to persist – means that high CN may be better at maintaining alleles that confer activity in environments that were not the target evolutionary pressure, leading to a form of evolutionary robustness.

**Figure 5.**
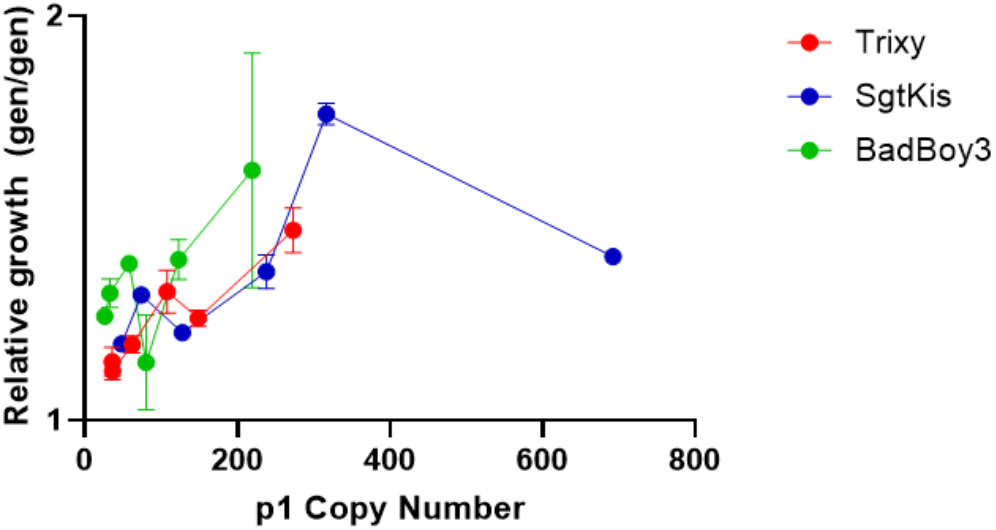
Comparison of relative activity of active allele libraries in acetamide versus butyramide. Variant libraries constructed for measuring fitness were grown for one passage in high concentrations of butyramide to reduce frequency of non-functional variants and then inoculated at equivalent OD_600_ into either 1 mM acetamide or 2 mM butyramide. After two days of growth, OD_600_ was measured and the number of doublings (generations) in either medium was determined. Plotted is the (number of generations reached in acetamide)/(number of generations reached in butyramide). On average, the variants produced by evolution under high CN had greater relative activity for acetamide over butyramide.

## Discussion

In this work, we developed OrthoRep into a versatile platform for exploring the evolutionary dynamics of multicopy plasmid systems. This required a strategy for changing the CN of the orthogonal p1 plasmid, which we achieved by expressing error-prone OrthoRep DNAPs at different levels using a series of yeast genomic promoters. In doing so, we were able to modulate p1’s CN over a ∼10-fold range for all three error-prone DNAPs tested. Notably, CNs stayed constant over at least 100 generations even when we subjected a p1-encoded gene to evolution for improved function.

The ability to set the CN and mutation rate of p1 allowed us to study how CN influences the adaptation of a model gene, amdSYM, encoded on p1. By comparing the evolution of amdSYM to improve its activity in utilizing butyramide as a sole nitrogen source at different CNs, we confirmed previous predictions on the evolutionary dynamics of multicopy plasmids. For example, we observed that high CN can act to reduce the exposure of deleterious mutations to purifying selection, resulting in the accumulation of a greater fraction of non-functional alleles. Likewise, we observed that high CN can act to dilute the fitness effect of beneficial mutations, resulting in their slower enrichment. The combination of these effects explains why continuous amdSYM evolution at low CN resulted in a higher fraction of functional alleles whose average fitness was also higher. Interestingly, the slower enrichment of amdSYM mutants specializing in butyramide metabolism in the high CN evolution experiments resulted in the greater maintenance of sets of alleles with higher activity on amdSYM’s original acetamide substrate, consistent with previous studies^11,14^ showing that multicopy plasmids support the coexistence of both ancestral and new traits during evolutionary innovation to favor robustness in more than one environment. Future work will be necessary to explore how far this type of tradeoff can be generalized. It is possible that while low CN ensures rapid traversal of greedy mutational trajectories, high CN maintains more sequence diversity during search in favor of robustness to more environments or increasing the navigability of rugged fitness landscapes.

A perhaps surprising observation from our experiments was that the distributions of fitnesses of evolved alleles were bimodal, consisting of a low mode of non-functional sequences and a high mode of functional sequences that could sustain butyramide metabolism in cells. In retrospect, this can be explained by a model wherein a given cell could afford to generate a substantial number of non-functional amdSYMs by mutation without suffering a fitness cost, because the aggregate activity of WT amdSYMs at the beginning of evolution or evolved amdSYMs at some point during evolution exceeded the gene dosage needed to maintain high fitness. Mutation of functional copies, which can easily lead to non-functional alleles since random mutations are mostly deleterious^30–32^, could then occur. Such a model is consistent with the fact that we found the low mode to be more populated with alleles when evolved under high CN conditions versus low CN conditions. An interesting consequence of this dynamic is that although we observed that low CN resulted in faster enrichment of certain high fitness alleles, this effect may be temporary, since evolution under high CN could eventually reach what would effectively be evolution under low CN once the non-functional fraction increases. In fact, this may be why in certain high CN evolution experiments where the growth rate of cells at the beginning of the evolution campaign was already high – suggesting little pressure to improve amdSYM – we still saw improved amdSYM alleles. Here, the idea is that some copies mutated to become non-functional, perhaps overshooting, resulting in an increase in the selection pressure on functional copies to adapt. Overall, the empirical observation that different fractions of copies became non-functional means that the effective CN of functional sequences that selection acted on was a dynamic parameter throughout evolution, perhaps itself evolving. Future work will be necessary to explore how the relationship between gene dosage and fitness, along with other variables such as mutation rate and mutation preferences, governs the effective functional CN reached in the evolution of multicopy plasmids. It is worth noting that the functional CN appears to plateau at high CNs, with the functional CN varying strongly based on the mutation rate (Fig. S7B). This suggests a complex relationship between gene dosage, segregation, and error rate which will be of interest to researchers studying natural high error rate systems.^33,34^

Our work also provides both an expanded toolkit and guidance in applying OrthoRep for directed evolution. For example, the observation that low CN promotes the faster enrichment of beneficial mutations and results in higher average fitness of functional alleles should favor the use of low CN systems in most applications. A caveat is that the fittest amdSYM alleles discovered were not necessarily from the lowest CN conditions, as the handful of alleles with the highest enrichment scores seemed comparable across different CN conditions (Fig. 4H-J). However, due to the precision needed to make comparisons on such few alleles from a high-throughput enrichment assay, we do not attempt to do so here.

This study does have some limitations. For example, the model gene we used to study plasmid evolutionary dynamics, amdSYM, did not need to adapt much (or at all) to achieve high butyramide activity for the host cell at many of the gene dosages represented across our CNs. Whether our study’s conclusions generalize to cases where large increases in activity and fitness are being evolved, as would pertain to neofunctionalization, antibiotic resistance, and most directed evolution applications, will require additional studies. Additionally, we assume that p1 randomly partitions during cell division, given a lack of annotation of an active segregation system^35^, but we do not know with certainty. Therefore, the relevance of our observations to multicopy plasmids with different segregation mechanisms remains unknown. Finally, we carried out all our evolution experiments in this work at a single somewhat small population size of 50,000 cells during passaging. It is plausible that larger population sizes would alter the distribution of outcome fitnesses observed and associated dynamics, for example by accessing and maintaining greater sequence diversity throughout evolution. Further studies addressing these limitations and exploring the evolutionary dynamics of multicopy plasmids with greater precision and versatility in general are now possible with our OrthoRep-based platform.

## Author Contributions

A.P. and C.C.L. conceived the project ideas. A.P. designed and carried out all experiments, collected and processed all data, and analyzed all data with input from C.C.L. A.P. and C.C.L. wrote the manuscript.

## Acknowledgements

This work was funded by NIH R35GM136297 to C.C.L. A.P. is supported by a Paul & Daisy Soros Fellowship for New Americans.

