## Supplementary Information for "A System to Explore the Adaptive Dynamics of Multicopy Plasmids: The Role of Copy Number and Mutation Rate in Evolutionary Outcomes"

**Supplementary Figures**

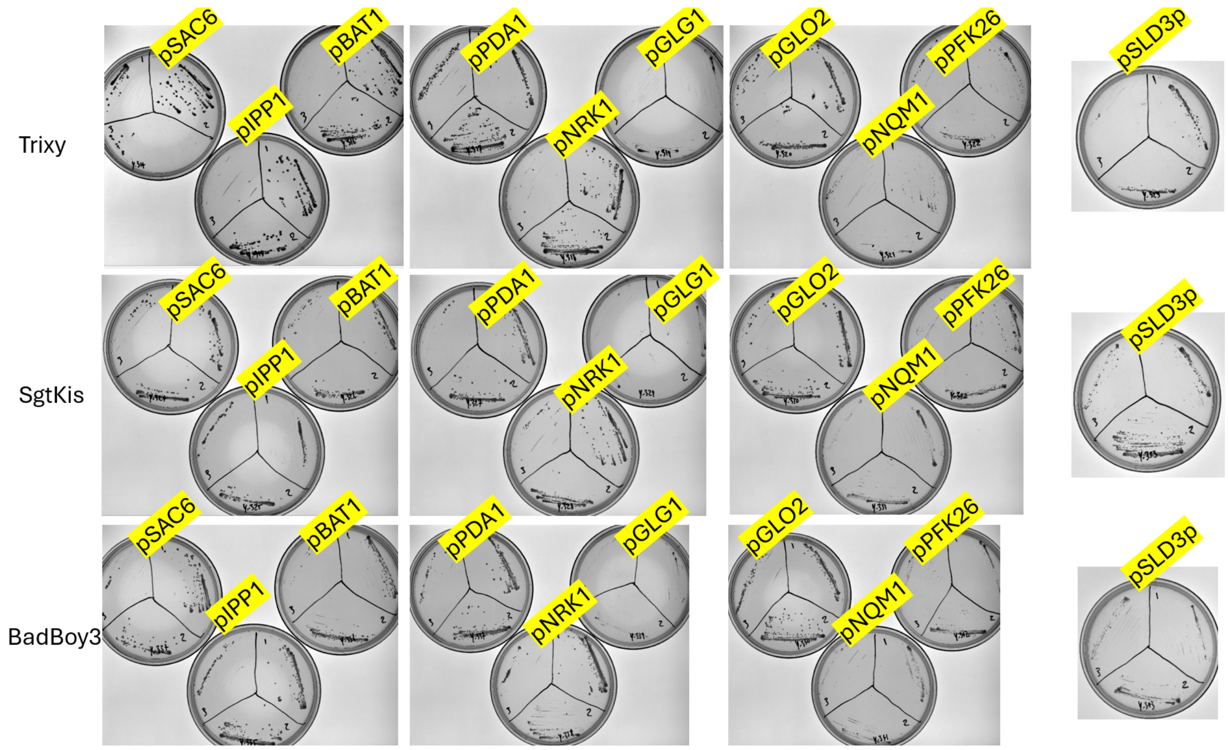


*Figure S1. Colony growth in relation to polymerase promoter strength.*

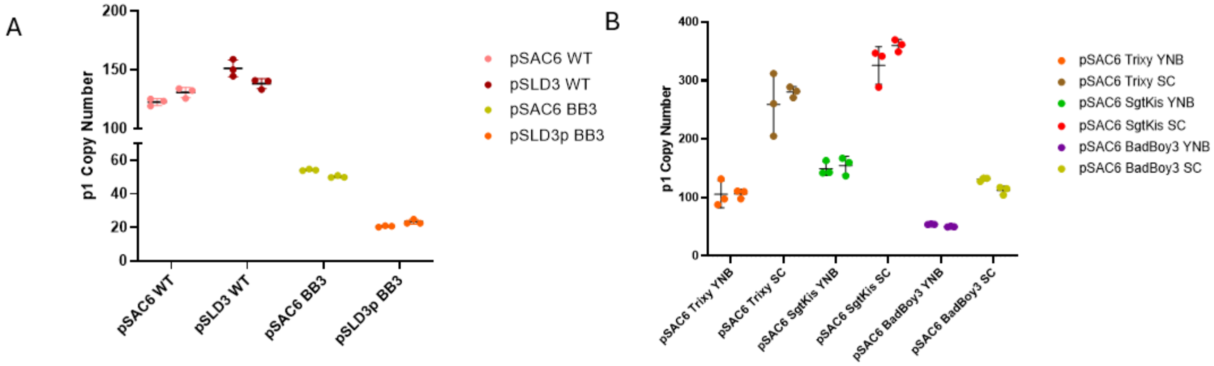


*Figure S2. CN relationships for error-prone vs wild-type polymerase and media conditions. A) Comparison between p1 copy number achieved from different expression strength of either an error-prone DNAP (BadBoy3 – BB3) or the WT low error rate DNAP. B) Comparison of p1 copy numbers achieved by pSAC6-driven DNAPs in two media conditions (SC, and minimal YNB media).*

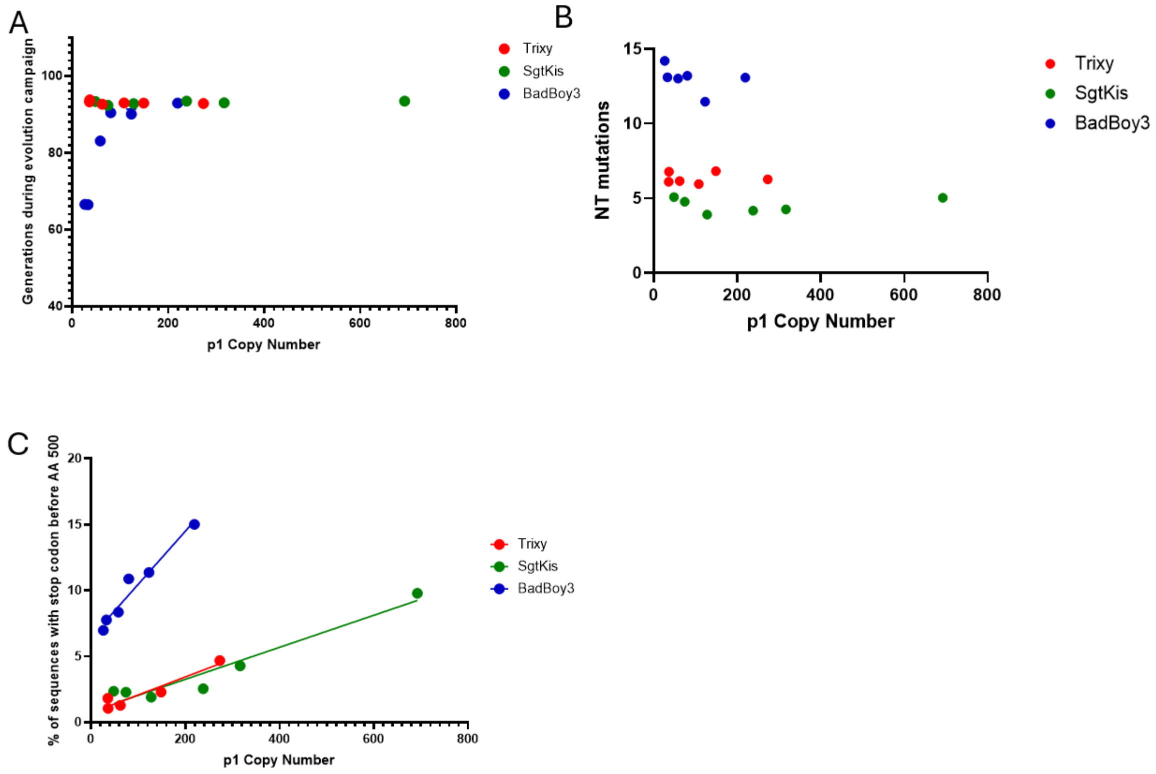


*Figure S3. Evolution metrics. A) The total number of generations of evolution at the given copy numbers and mutation rates. B) Mean number of nucleotide (NT) mutations per sequence as a function of copy number. There is no obvious correlation across different copy numbers. C) Analysis of fraction of sequences containing stop codons before amino acid 500 as a proxy for function, from nanopore sequencing. Higher copy is well correlated with more stop codon containing sequences, indicating more hitchhiking by non-functional mutants.*

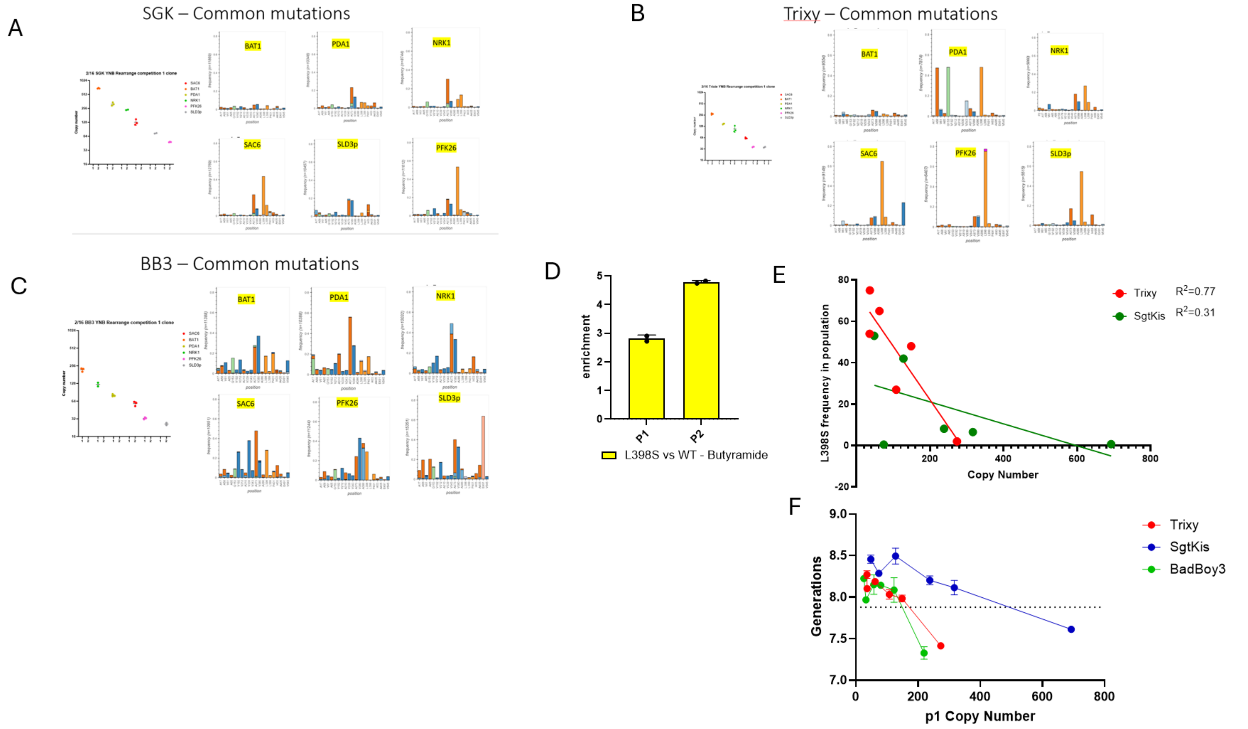


*Figure S4. Analysis of mutation frequencies and L398S. A-C) MAPLE analysis of mutation frequencies of amdSYM with specific attention paid to mutation L398S for SgtKis (A), for Trixy (B), and for BadBoy3 (C). D) Competition between L398S amdSYM variant and WT amdSYM. Variant L389S was transformed into a green fluorescent strain and WT amdSYM was transformed into a red fluorescent and competition was carried out on the mixed population. E) Frequency of L398S mutation plotted against CN during evolution. Note that one data point (pSLD3) is responsible for relatively decreased $R^2$ value for SgtKis, due to its discovery of other beneficial mutations. Omission of that data point would increase $R^2$ to 0.64. F) Number of generations reached during growth of bulk culture (two replicates) in the competition assay after 48 hours. Dotted line corresponds to the number of generations a WT amdSYM only control reached under the same growth conditions. Greater growth for the lower CN conditions suggests that the pool of evolved alleles from those conditions are more fit on average. Note, the pNRK1-driven Trixy condition was nanopore sequenced separately. Results of nanopore from separate sequencing are shown here. Both enrichment data and separate nanopore data are still usable individually, and this does not affect any claims made in the paper. Variants with UMI's used for enrichment analysis, and the enrichment data, can be found in Table S3.*

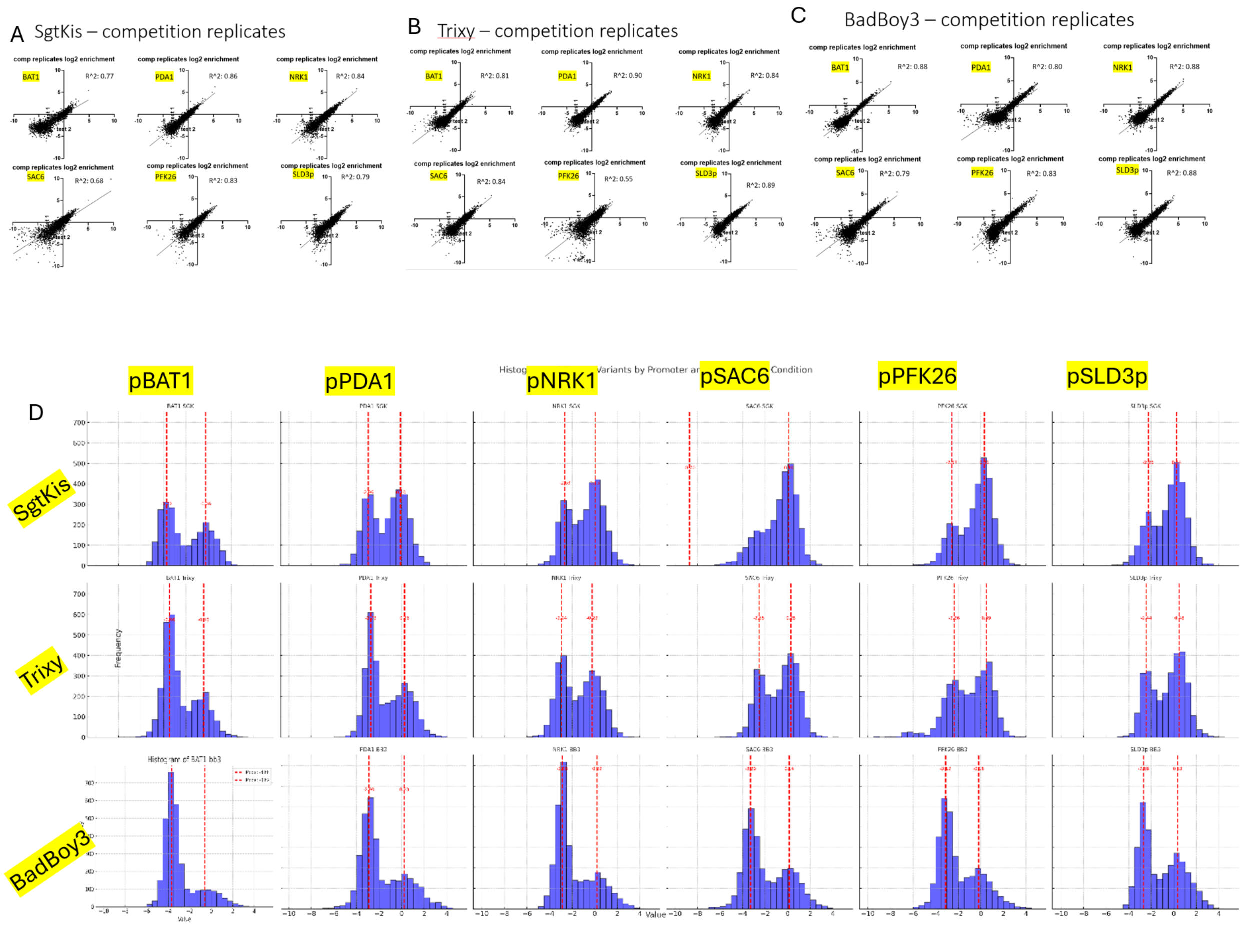


*Figure S5. Enrichment data for competition assays. A-C) Correlation plots for competition assay replicates for SgtKis (A), Trixy (B), and BadBoy3 (C). D) Data reduced to histograms for modal calculation.*

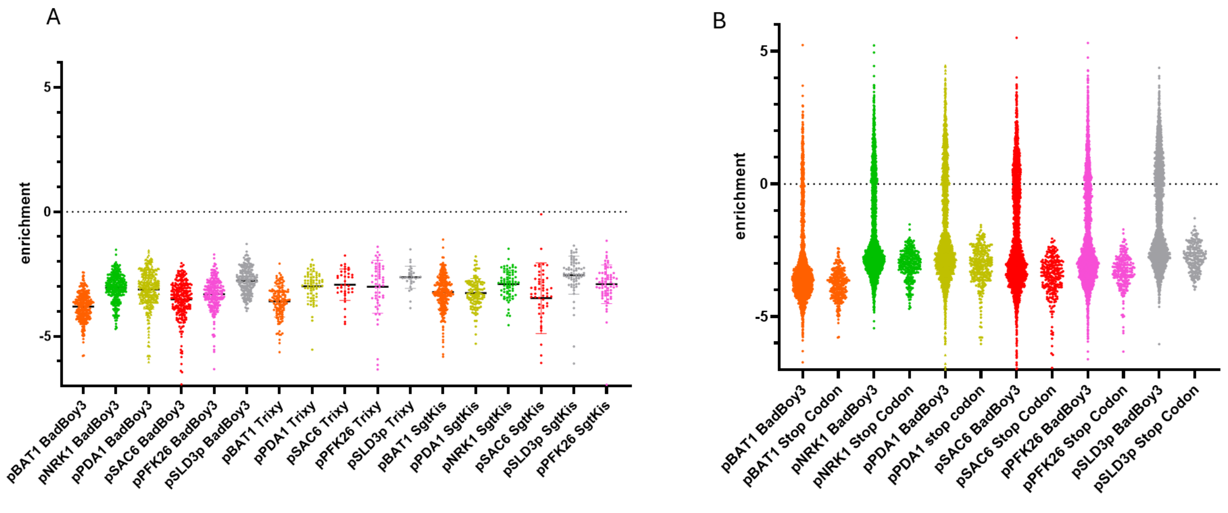


*Figure S6. Enrichment data for stop codon containing sequences. A) Competition data filtered for only sequences that contain stop codons prior to amdSYM amino acid 500 to outline the enrichment factor range of non-functional sequences. B) Stop codon only sequences next to all variants of BadBoy3 samples to show that all sequences in the low mode are equivalent to non-functional sequences.*

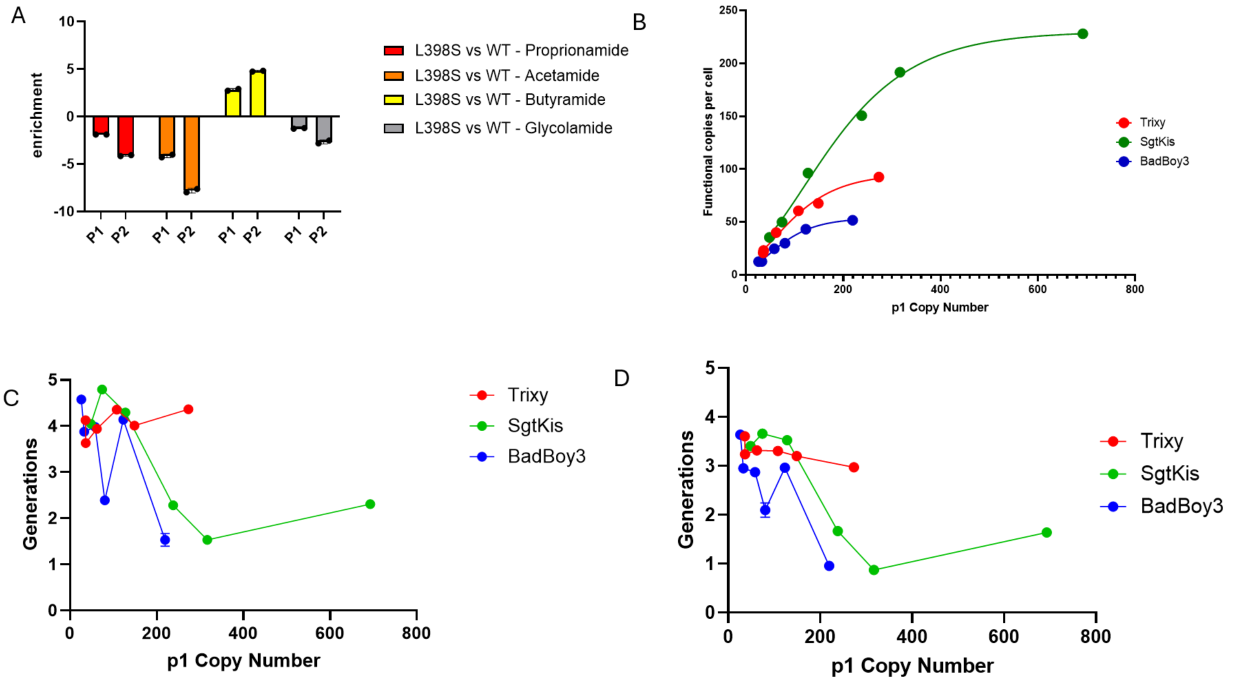


*Figure S7. Evaluation of specialization during evolution. A) Evaluation of the effect of L398S on activity for proprionamide, acetamide, butyramide, and glycolamide. L398S increases activity on butyramide but decreases activity on 3 other amides. B) Functional copies per cell, calculated by multiplying the proportion of variants in the high mode (see Fig. 4H-J) by the CN at the final evolution timepoint. C-D) Growth of evolved and functionally enriched allele sets in acetamide (C) and butyramide (D). To calculate relative activity shown in Fig. 5, the generations reached in acetamide in 48 hours of growth was divided by the generations reached in butyramide in 48 hours of growth. We note that although the high CN conditions had lower growth rate (fewer generations reached in 48 hour) in both acetamide and butyramide in this assay, these growth rates are measured for populations where each cell only encodes one copy of the amdSYM allele. During the evolution campaign itself, each cell contains multiple amdSYM allele copies and the high CN cases have a higher number of function copies than the low CN cases (B). Given this and the fact that in the low CN butyramide evolution experiments, the evolved growth rate of cells was mostly equivalent to the growth rate of cells in the high CN evolution experiments (Fig. S3A), we conclude that the high CN condition in the actual multicopy plasmid evolution experiment context would exhibit equal growth rate in the butyramide condition as the low CN conditions even though the high CN conditions in this assay that measures per allele average fitness had lower growth rate. Thus, the higher average activity of alleles from the high CN conditions for acetamide relative to butyramide compared to the low CN conditions means that in the multicopy plasmid evolution experiment, the cells from the high CN condition retained more of their plasmid-encoded ancestral acetamide activity than cells from the low CN condition.*

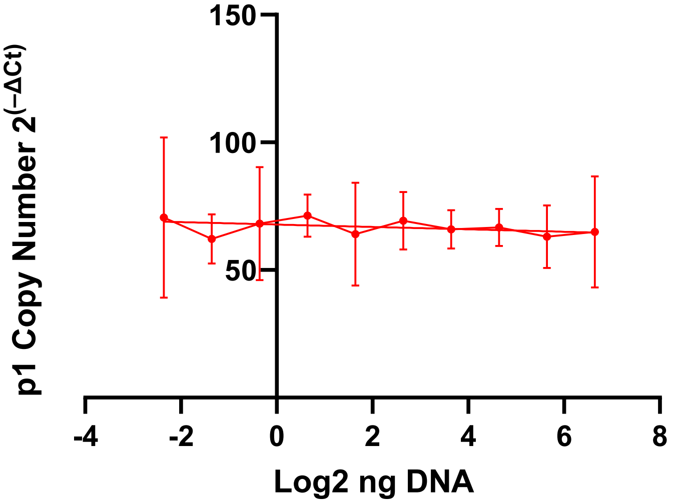


*Figure S8. Copy number standard curve. A copy number standard curve was generated for each qPCR run by serial 1:2 dilution of one template. If primer efficiency varied between genomic normalization gene (Gal1) and p1 copy number gene (Leu2), the apparent copy number would change across the serial dilution when using the $2^{(-\Delta Ct)}$ method.*

### Materials and Methods

#### DNA plasmid construction

All plasmids used in this study are listed in Table S2. DNA templates for PCR were obtained from previously published constructs or ordered as gBlocks (Integrated DNA Technologies, IDT). Oligonucleotide primers were purchased from IDT. PCR reactions were performed using PrimeSTAR GXL DNA polymerase (Takara, Cat. No. R050A) under standard conditions recommended by the manufacturer. Assembled plasmids were generated via Gibson Assembly or Golden Gate Assembly (New England Biolabs, NEB), following the manufacturers' protocols. Chemically competent *Escherichia coli* TOP10 cells (ThermoFisher) were transformed with the assembled products and cultured using standard conditions. Promoter regions for polymerases were amplified from *Saccharomyces cerevisiae* genomic DNA. Resulting plasmids were sequence-verified by whole-plasmid sequencing (Plasmidsaurus). Detailed information on all plasmids and primers, including those used to amplify genomic promoters, can be found in Table S2.

#### Yeast culture

The base yeast strain was *S. cerevisiae* BY4741/4742[1]. A MetP1 landing pad was introduced by protoplast fusion[2]. Yeast were routinely cultured in Synthetic Complete (SC) media composed of Yeast Nitrogen Base (YNB) + amino acid/supplement mix (A/S) and 2% glucose. Selection on antibiotic-containing media was performed using YPD agar supplemented with Hygromycin B, Nourseothricin (NAT), or G418 at appropriate concentrations. When antibiotics were combined with dropout media, YNB + A/S was replaced by YNB – A/S plus 1 g/L monosodium glutamate (Sigma-Aldrich). Butyramide and other amides (Sigma-Aldrich) were prepared as 400 mM stocks in sterile-filtered distilled water.

#### Evolution campaign

For the evolution experiments, yeast were passaged in YNB media lacking amino acids and A/S (US Bio) with 2% glucose and an initial concentration of 4 mM butyramide. The yeast strains used were engineered to become prototrophic upon receiving AmdSym-Leu2 P1. Amino acids were omitted because the yeast can use them as nitrogen sources. At passage 7 (out of 13), the butyramide concentration was reduced to 2 mM. Optical density (OD600) was measured at each passage, and cultures were normalized to transfer approximately 50,000 cells into 1 mL fresh media (48-well block format). Cultures were passaged daily and never allowed to reach saturation.

#### Yeast transformation / p1 transfer

Transformations were carried out using a previously described frozen competent cell protocol[3]. Briefly, cells were grown overnight in selective media, back-diluted the following day to OD600 = 0.1, and then grown in YPD to OD600 = 1.2. Cells were washed twice in water and frozen at –80°C in a solution of 10% DMSO and 5% glycerol. Transformations were performed by mixing approximately 54 µL of DNA solution with 260 µL PEG, 36 µL LiAc, and 10 µL salmon sperm ssDNA (ThermoFisher), followed by heat shock. EcoRI-linearized promoter-polymerase plasmids were integrated into strain y_256, which carries a genomic I-SceI site for efficient integration (OP_202). For the NRK1 gene, which contains an internal EcoRI site, BglII digestion was used. Transformants were selected on YPD + NAT plates, and successful integration was confirmed by screening for NAT resistance, loss of Hygromycin resistance, and by outside-outside PCR.

For P1 integration, yeast *y_215* (Table S2) carrying a genomically encoded wild-type polymerase (low error rate) and a MetP1 landing pad (GR419) were transformed with ScaI-digested P1 integration constructs. Cells were passaged 5x times in -L and integration was confirmed by yeast miniprep[4] and agarose gel analysis. P1 transfer to a new strain was achieved via abortive mating[5]. Equal numbers of donor and recipient cells were mixed, plated on YPD for 6 hours, then restreaked on –HLWMCR NAT/CAN plates. After 3 days, colonies were picked into SC-HLUWMC media. Following 24 hours of growth, cells were passaged in parallel into minimal media and SC-HLUWMC for copy number measurements.

**qPCR copy number quantification**

Genomic DNA was prepared by GC prep[6]. Yeast OD600 was measured (Tecan Spark), and cells were normalized to 1 mL at OD600 = 0.4. Pelleted cells (17,000 × g, 1 min) were resuspended in 100 µL 5% Chelex (Sigma) with acid-washed glass beads and disrupted by vortexing for 6 min. Samples were then boiled at 100°C for 6 min, centrifuged, and the supernatant used as template DNA.

qPCR reactions (10 µL final volume) included 5 µL of 2× SYBR PowerUp mix (Thermo, Cat. No. A25742), 0.04 µL of 100 µM primers, 2 µL ddH2O, and 3 µL of a 1:10 dilution of genomic DNA. Reactions were run in white 384-well plates (Thermo, Cat. No. 164610) with a standard (not fast) SYBR protocol for 40 cycles on a QuantStudio 6 (Applied Biosystems).

To ensure equal primer efficiency, a control sample was serially diluted. Primers targeted GAL1 (genomic control) and LEU2 (on P1). Because the host lacks LEU2 in the genome, all LEU2 signal originated from P1. Each sample was run in biological duplicates and technical triplicates. Copy number was calculated as $2^{(-\Delta Ct)}$, where $\Delta Ct = Ct(LEU2) - Ct(GAL1)$. The instrument was configured to treat primers as having a single target to avoid automatic threshold adjustments. Given equal primer efficiency (Supplemental Fig. 8), efficiency corrections were not required. All primers are listed in Table S2.

**Growth curves**

Yeast containing amdSYM-encoded P1 were passaged twice in YNB + glucose + 4 mM butyramide to deplete excess nitrogen stores. On the measurement day, cells were normalized to OD600 = 10 in minimal media, then diluted 1:100 into 200 µL YNB + glucose + ammonia media in microplate wells. Cells were grown at 30°C with continuous shaking, and OD600 was recorded every 15 min. Growth curves were performed in biological duplicate. The growth rate was calculated based on the minimum doubling time.

**Flow cytometry / measuring expression**

For expression analysis, an mKate-Leu P1 (generated with OP_096, table S2) was transferred into recipient cells containing polymerases driven by BAT1, SAC6, PFK26, or SLD3 promoters. After mating and selection (as described above), two replicate colonies were picked into –L media and grown for 24 hours. Flow cytometry was performed using a ThermoFisher Attune cytometer. Cell gating ensured size and single-cell events. The geometric mean fluorescence intensity was recorded. Passaging was minimized to prevent confounding by variable error rates.

**PCR assembly/UMI tagging**

Variants of the amdSYM gene were amplified from evolution samples using genomic DNA (prepared as for qPCR). Primers included the first 8 amino acids of the amdSYM gene and a portion of P1 downstream of the gene to capture 3'-end mutations. PCR was performed with Platinum SuperFi 2× mix (Invitrogen) with minimal cycles. Amplicons were gel-extracted.

Two genomic integration flanks were PCR-amplified from a plasmid (OP_209). One flank contained a TP901 AttP site and the POP6 promoter; the other included a unique molecular identifier (UMI), a LEU2 marker, and another TP901 AttP site. The UMI was introduced using a PAGE-purified primer with a 20× NNN degenerate sequence. PCR conditions were optimized for minimal cycling, and products were DpnI-treated and gel-extracted. Assembly PCR included 5 ng of each amplicon (gene library + flanks) and only external primers, and was performed with GXL polymerase (Takara). A sample cassette is listed in Table S2 (OP_comp).

**Library Integration / down sampling**

Approximately 500 ng of PCR-assembled amplicons were transformed into yeast containing a genomic TP901 landing pad at the HO locus (OP_205) and a 2 µm plasmid expressing TP901 recombinase (OP_189). Competent cells were prepared using the frozen competent cell protocol. Transformation efficiency was determined by plating 1/1000 of the transformed culture. After two days, a fraction of the plate corresponding to ~10,000 sequences per library was scraped and banked. A wild-type (WT) control was prepared separately using a plasmid (OP_209) template for amdSYM, with the construct/UMI sequenced separately.

**Nanopore**

Transformed libraries were grown, and DNA was prepared by GC prep. Primers flanking the genomic locus and the UMI were used to amplify the cassette with SuperFi polymerase, minimizing cycle number. Amplicons were gel-extracted and further barcoded by a 10-cycle PCR using a distinct downstream primer with a 7 bp barcode. Amplicons were ExoSAP-treated, column-purified, quantified by Qubit HS (Thermo), normalized, and pooled. Libraries were prepared with the LSK-114 Nanopore kit and sequenced on a single flow cell. Analysis and demultiplexing were performed using the MAPLE pipeline (https://github.com/gordonrix/maple).

**Competition assay and Illumina sequencing**

Libraries were passaged twice in nitrogen-limited media to deplete internal nitrogen stores. The WT control was spiked in at a 1:500 ratio by OD. For each library, ~200,000 cells were inoculated into 2 mL YNB + 2% glucose + 2 mM butyramide in duplicate wells. A 1 mM ammonia control was included separately. Day 0 DNA samples (inoculum) were prepared by GC prep.

After 48 hours of growth, cultures were harvested. OD600 measurements ensured normalization of cell density. DNA was prepared via GC prep. The UMI region was amplified from ~5 ng DNA with minimal cycles, confirmed on a 2% agarose gel, and treated with ExoSAP-IT (Thermo Fisher). A second PCR added barcode handles and Illumina P5/P7 adapters. Samples were sequenced on half of a NovaSeq X lane and demultiplexed by Azenta.

UMI counts were normalized to WT, and enrichment scores were calculated as:

$$Enrichment = log2\frac{(\%\,Evolved/\%Control)T_x}{(\%\,Evolved/\%Control)T_0}$$

Only the top 3000 sequences (by abundance at T=0) were considered. At T=0, the 3000th UMI had >1000 reads. Mean values of biological replicates were used for analysis and plotting.

**Relative activity of libraries on acetamide and butyramide**

Because weakly expressed pPOP6-driven amdSYM variants grew too slowly for reliable growth curves, samples were prepassaged in 2 mM ammonia and then 60 mM butyramide media. Cells were normalized to OD600 = 5, then inoculated at OD600 = 0.05 into either 2 mM acetamide or 4 mM butyramide media in duplicate. After 2 days, OD600 was measured, and generations were calculated as log2(OD/0.05). The ratio of generations in acetamide to generations in butyramide was reported.

**Comparison of L398S activity vs. WT across substrates**

An L398S amdSYM variant, driven by the POP6 promoter in a similar cassette as the pooled competition but lacking a UMI, was transformed into a green-fluorescent strain (y_081, Table S2). WT amdSYM was introduced into a red-fluorescent strain (y_082, Table S2). Colonies were grown in SC-HLUWMC and then passaged once in minimal YNB + 4 mM ammonia. WT and L398S strains were mixed and inoculated 1:50 into minimal YNB containing either 2 mM propionamide, 1 mM acetamide, 4 mM butyramide, or 16 mM glycolamide. Initial cell ratio was measured by flow cytometry at day 0. After 24 and 48 hours, ratios were measured again. Log2 enrichment was calculated as above.

**Competition assay analysis**

<u>Data Acquisition and Preprocessing:</u>

Paired-end sequencing data were generated using the Illumina NovaSeq platform, resulting in 72 unique molecular identifier (UMI) libraries. The data were demultiplexed to produce individual FASTQ files for each sample. Forward and reverse reads were stored in files named following the convention sample_R1_001.fastq and sample_R2_001.fastq.

<u>Read Merging</u>

Overlapping paired-end reads were merged to reconstruct the full-length amplicons using BBMerge (version 39.08) from the BBTools suite. Merging was performed to improve the accuracy of downstream analyses by combining overlapping read pairs into single sequences. The merging process was executed with the following command for each sample:

```
bash
bbmerge.sh in1=sample_R1_001.fastq in2=sample_R2_001.fastq
out=merged_sample.fastq
```

This command produced merged FASTQ files (merged_sample.fastq) containing the combined sequences of forward and reverse reads.

<u>Reference Sequence Preparation</u>

A reference sequence was constructed to target the specific variable region within the amplicon. The region of interest consisted of a 20-base variable sequence flanked by constant regions. The reference sequence was defined as:

shell

```
>target_region
TAGGGCNNNNNNNNNNNNNNNNNNNNGAGAGA
```

This sequence was saved in FASTA format as reference.fa and used for alignment.

<u>Alignment</u>

Merged reads were aligned to the reference sequence using Bowtie2 (version 2.4.4) in very-sensitive mode to maximize alignment sensitivity and accuracy. The reference index was built using the following command:

bash

```
bowtie2-build reference.fa reference_index
```

Alignment of the merged reads to the indexed reference was performed with:

bash

```
bowtie2 -x reference_index -U merged_sample.fastq -S aligned_sample.sam --very-sensitive
```

This generated SAM files (aligned_sample.sam) containing the alignment results for each sample.

<u>Variable Region Extraction and Counting</u>

A custom Python script was developed to extract the 20-base variable region from the aligned reads and count the occurrences of each unique sequence. The script performed the following steps:

<u>Parsing SAM Files:</u> The script read the SAM files and extracted the read sequences aligned to the reference.

<u>Identifying Variable Region:</u> Using the known position of the variable region within the reference sequence, the script calculated the start and end indices corresponding to the variable region in each read.

<u>Extracting Variable Sequences:</u> The script extracted the 20-base variable region from each read sequence based on the calculated indices.

<u>Filtering and Counting:</u> Only sequences of the correct length (20 bases) without ambiguous nucleotides were considered valid. The script counted the occurrences of each unique variable sequence.

<u>Outputting Results:</u> The counts were saved in a CSV file (variable_region_counts.csv) with two columns: Variable Region and Count.

<u>UMI Counting</u>

The unique molecular identifiers (UMIs) present in the variable region were effectively counted through the extraction and counting process described above. Each unique sequence in the variable region corresponded to a distinct UMI, allowing for quantification of UMI diversity and abundance within the samples.

<u>Software and Computational Environment</u>

<u>BBTools (BBMerge) Version 39.08:</u> Used for merging paired-end reads.

<u>Bowtie2 Version 2.4.4</u>**:** Utilized for aligning merged reads to the reference sequence.

Python 3.8: Employed for scripting and data processing, with the pandas library for handling data structures.
Operating System: Ubuntu 20.04 LTS running under Windows Subsystem for Linux (WSL 2).
Command Summary
Read Merging:
bash
bbmerge.sh in1=sample_R1_001.fastq in2=sample_R2_001.fastq
out=merged_sample.fastq
Reference Indexing:
bash
bowtie2-build reference.fa reference_index
Alignment:
bash
bowtie2 -x reference_index -U merged_sample.fastq -S aligned_sample.sam --very-sensitive

Variable Region Counting (Python Script Overview):
Input**:** aligned_sample.sam
Process: Extract variable region based on known indices, count unique sequences.
Output: variable_region_counts.csv
Data Availability
All scripts and command-line tools used in this analysis are available upon request. The custom Python script for variable region extraction and counting can be provided to facilitate reproducibility.
Generating histograms and analyzing modes:
To assess the distribution and identify the most common values (modes) of different fitness variants obtained during the evolution experiments, we conducted a detailed analysis using histograms and kernel density estimates.
Data Processing: The dataset comprised multiple columns representing fitness values for distinct evolutionary conditions. Each column was analyzed separately to determine its distribution. Missing values were removed prior to analysis.
Kernel Density Estimation (KDE**):** To identify the two most prominent modes (peaks) for each fitness variant, we applied Gaussian Kernel Density Estimation (KDE). KDE was used to estimate the underlying probability density function of each column's values, generating a smoothed representation of the data distribution. Local maxima were then identified as candidate modes.
Mode Selection: For each fitness column, the two most prominent modes were selected based on the density estimates. When fewer than two distinct local maxima were present, the existing maxima were noted, and the absence of additional modes was reported.
Histogram Generation: Histograms were generated to visualize the distribution of values for each fitness variant. Bins were defined to adequately represent the spread of the data. The two identified modes were marked on each histogram using red dashed lines to facilitate visual interpretation.

Visualization and Export: Histograms for all fitness variants were generated and saved as individual PNG files. These visualizations were then compiled into a single ZIP archive for easy reference and further analysis.

Variable Region Proportion Analysis

A Python script was developed to process and analyze variable regions from sequencing data across four experimental conditions: (1) Day Zero, (2) Ammonia Stress, (3) Butyramide Replicate 1, and (4) Butyramide Replicate 2. The input data were CSV files containing counts of unique variable region sequences.

1. Data Preprocessing:
   - CSV files for each condition were read into Python using pandas.
   - The proportion of each variable region within its condition was calculated by dividing its count by the total read count for that file.
2. Wild-Type Normalization:
   - A predefined WT sequence (GCGCGTCTTATACCGGAACA) was used as a reference.
   - For each sequence, its relative proportion compared to the WT was calculated.
3. Top Sequence Selection:
   - From the Day Zero dataset, the top 3,000 sequences (by abundance) were identified.
4. Cross-Condition Comparison:
   - The proportions of these top sequences from Day Zero were matched across all conditions.
   - Missing data for sequences not present in certain conditions were replaced with a value of 0.
5. Output:
   - The final data table included each variable region and its relative proportions across conditions, saved as a CSV file (BAT1BB3_analysis_output.csv).
6. Execution Environment:
   - The script was executed in a Windows Subsystem for Linux (WSL) environment to facilitate file path compatibility.
7. Outputs:
   - Results were stored in an output directory for subsequent analysis.
